# Mice deficient of *Myc* super-enhancer region reveal a differential control mechanism between normal and pathological growth: *MYC* non-coding region regulating cancer growth

**DOI:** 10.1101/082602

**Authors:** Kashyap Dave, Inderpreet Sur, Jian Yan, Jilin Zhang, Eevi Kaasinen, Fan Zhong, Leander Blaas, Xiaoze Li, Robert Månsson, Jussi Taipale

## Abstract

The gene desert upstream of the *Myc* oncogene on chromosome 8q24 contains susceptibility loci for several major forms of human cancer, including cancers of breast, prostate, and colon. The region shows high conservation between human and mouse and contains multiple *MYC* enhancers that are activated in tumor cells. However, the role of this region in normal development has not been addressed. Here we show that a 538 kb deletion of the entire *MYC* upstream super-enhancer region in mice results in 50 to 80% decrease in *MYC* expression in multiple tissues. The mice are viable and show no overt phenotype. However, they are resistant to tumorigenesis, and most normal cells isolated from them grow slowly in culture. Consistently, deletion of the 8q24 super-enhancer region perturbs Myc targets only in cultured cells, but not *in vivo*. These results reveal that only cells whose Myc activity is increased by serum or oncogenic driver mutations depend on the 8q24 super-enhancer region, and indicate that targeting the activity of this element is a promising strategy of cancer chemoprevention and therapy.

## Introduction

Deregulated expression of the *MYC* oncogene is associated with many cancer types (Reviewed in (Albihn et al. 2010; Dang 2012; Evan 2012)). MYC acts primarily as a transcriptional activator that increases expression of many genes required for RNA and protein synthesis above the level that is required in resting cells. In cancer cells, aberrantly elevated levels of MYC drive global amplification of transcription rates, providing the cells with necessary resources for rapid proliferation (see, for example (Brown et al. 2008; van Riggelen et al. 2010; Ji et al. 2011; Lin et al. 2012; Sabo et al. 2014; Walz et al. 2014)).

Transcription of the *MYC* gene is regulated by a diverse array of regulatory elements located both upstream and downstream of the *MYC* transcription start site (TSS). Variants in the *MYC* upstream region contribute to inherited susceptibility to most major forms of human cancer, and account for a very large number of cancer cases at the population level (Amundadottir et al. 2006; Gudmundsson et al. 2007; Yeager et al. 2007; Al Olama et al. 2009; Yeager et al. 2009). For example, the polymorphism rs6983267 linked to colorectal (Tomlinson et al. 2007) and prostate (Yeager et al. 2007) cancers contributes more to cancer morbidity and mortality than any other known inherited variant or mutation, including the inherited mutations in classic tumor suppressors such as *RB*, *TP53* and *APC*. Through computational and experimental analyses, we and others have shown that the risk allele G of rs6983267 creates a strong binding site for the colorectal-cancer associated transcription factor Tcf7l2 (Pomerantz et al. 2009; Tuupanen et al. 2009). This binding site is located within the Myc-335 enhancer element that is dispensable for mouse viability, but required for efficient Tcf7l2-driven intestinal tumorigenesis (Sur et al. 2012).

More recently, another enhancer element, located 1.47 Mb downstream of *Myc* was shown to be required for formation of acute lymphoblastic leukemia (ALL) in mice (Herranz et al. 2014). However, in contrast to the Myc-335 element, this element is also required for normal T-cell development. Thus, the mechanism by which individual Myc enhancer elements contribute to normal development and tumorigenesis is still unclear.

Several studies have shown that the 8q24 region contains a large number of additional enhancer elements (see, for example (Hallikas et al. 2006; Ahmadiyeh et al. 2010; Yan et al. 2013; Yao et al. 2014)), forming a ‘super-enhancer’ region that is active in many different types of human cancer (Hnisz et al. 2013; Loven et al. 2013; Zhou et al. 2015). The *MYC*-associated super-enhancers are activated during the process of carcinogenesis (Hnisz et al. 2013), and downregulation of super-enhancer activity leads to selective inhibition of *MYC* expression (Loven et al. 2013). Thus, *MYC*-associated super-enhancer activity is required for tumorigenesis, but the role of these elements in normal tissue morphogenesis and homeostasis has been unclear.

To address this problem, we have in this work generated multiple mouse strains deficient of regulatory elements upstream of the *Myc* promoter. We found that surprisingly, the entire super-enhancer region conferring multi-cancer susceptibility contributes to Myc expression *in vivo*, yet is not required for mouse embryonic development and viability. However, this region, is required for the growth of normal cells in culture and cancer cells *in vivo*, thus identifying a common mechanism of growth control between the two cellular states.

## Results

### Functional mapping of the super-enhancer region upstream of Myc

To dissect functional significance of the 8q24 region during normal development, we generated series of *Myc* alleles in mice using homologous recombination in ES cells. These include the Myc-335 enhancer deletion allele we have described previously (Sur et al. 2012), and deletions of two additional conserved enhancer elements, Myc-196 and Myc-540, both of which are active in mouse intestine and colorectal cancer cells. In addition, we generated a point mutation that inactivates a conserved CCCTC-Binding factor (CTCF) site 2 kb upstream of the *Myc* TSS. This site has previously been reported to be required for *MYC* expression (Gombert and Krumm 2009), and to have insulator activity (Gombert et al. 2003) (Fig. 1a). Each allele contained loxP site(s) in the same orientation to allow conditional knockouts of the enhancers, and to facilitate generation of large deletions and duplications by interallelic recombination (Wu et al. 2007). All alleles were bred to homozygosity, and resulted in generation of viable mice. Expression of *Myc* in the colon of Myc-196^−/-^ and Myc-540^−/-^ mice was not markedly altered, suggesting that these elements have little effect on regulation of *Myc* in the intestine under normal laboratory conditions (Fig. 1b). *Myc* expression level was also normal in Myc-CTCF^mut/mut^ mouse colon despite complete loss of CTCF binding to the region proximal to the *Myc* promoter (Fig. 1c).

**Figure 1:**
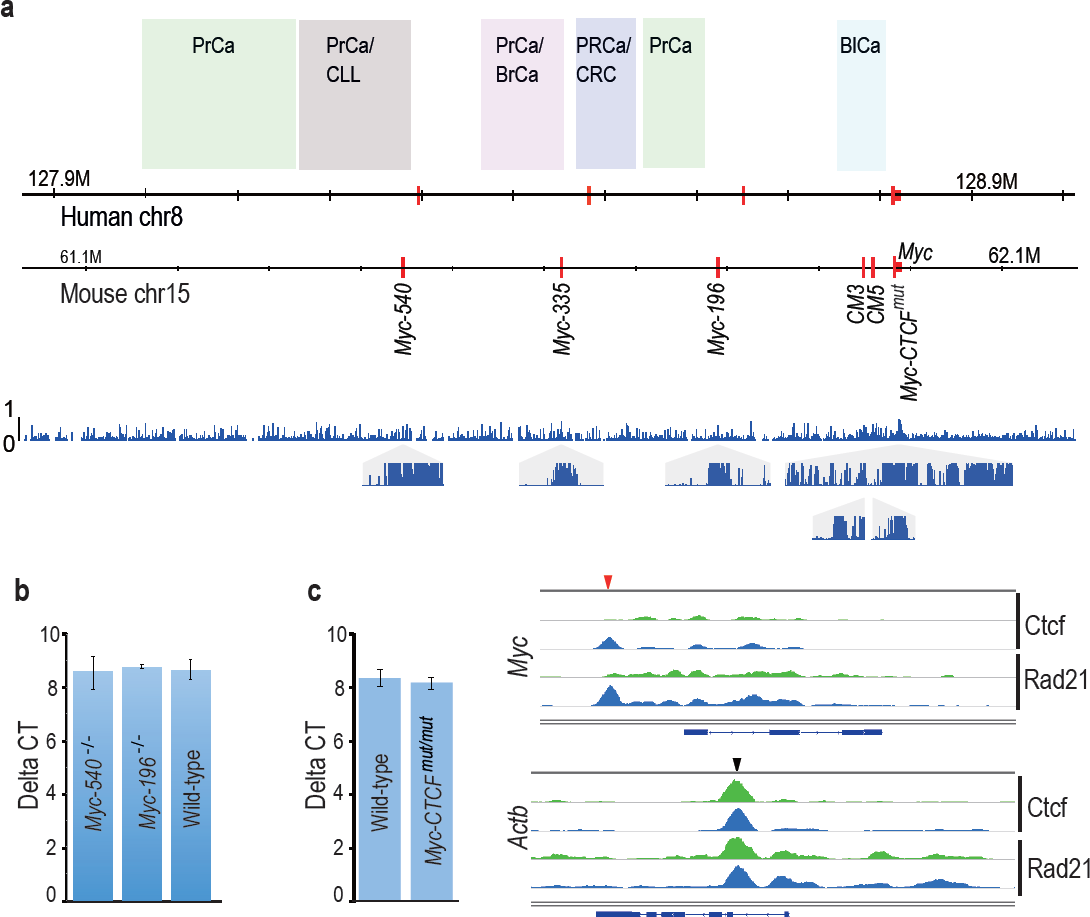
Cancer susceptibility region upstream of Myc contains several conserved enhancer elements that are dispensable for normal mouse development and Myc expression. (a) Comparison of Myc locus between human and mouse. The susceptibility regions for prostate cancer (PrCa), chronic lymphocytic leukemia (CLL), breast cancer (BrCa), colorectal cancer (CRC) and bladder cancer (BlCa) are marked. Red vertical lines mark the location of the Tcf7l2-binding colorectal Myc enhancers in the two species. The lower panel shows the regional conservation probability predicted by PhastCons (hg19 assembly, UCSC) with non-overlapping sliding windows for the whole region and each enhancer locus with a size of 500 bp and 10 bp, respectively. (b) Deletion of Myc-196 and Myc-540 enhancer elements does not affect Myc expression in the colon as determined by qPCR analysis (Myc-196^−/-^ *n*=2, Myc-540^−/-^ *n*=3 and wild-type *n*=5). (c) Mutation of the Myc-CTCF site causes loss of CTCF and Rad21 binding at the *Myc* locus (top panel). Binding of CTCF and Rad21 at a control *Actb* locus is not affected. Red and black arrowheads denote binding sites at *Myc* and *Actb* loci, respectively; green: Myc-CTCF^mut/mut^, blue: wild-type. The gene body for *Myc* and *Actb* is shown below the respective panels. The qPCR analysis reveals that despite loss of CTCF/cohesin binding, the expression of Myc mRNA is not altered in the colon (for qPCR, Myc-CTCF^mut/mut^ *n*=4, wild-type *n*=3). Error bars denote one standard deviation.

### Mice lacking the Myc super-enhancer region are viable and fertile

As the individual mutations and deletions had limited effect, we next decided to generate two large deletions in the *Myc* locus using interallelic recombination between the Myc-CTCF^mut^ loxP site and the loxP sites at Myc-335^−^ and Myc-540^−^, yielding deletions of 365 kb (GRCm38/mm10 chr15:61618287-61983375) and 538 kb (chr15:61445326-61983375), respectively (Fig. 2a). The resulting alleles, Myc^Δ^2-367^^ and Myc^Δ^2-540^^, were then segregated out from the corresponding duplications, and bred to homozygosity. Given the very large regions that were deleted (Fig. 2b), we expected to see a strong phenotype. However, no overt phenotype was identified in the Myc^Δ^2-367/^Δ^2-367^^ mice. The mice were born at the expected mendelian ratio, and both males and females were viable and fertile. Analysis of *Myc* expression, however, revealed a strong decrease in *Myc* expression in the colon and ileum of the mice (not shown).

**Figure 2:**
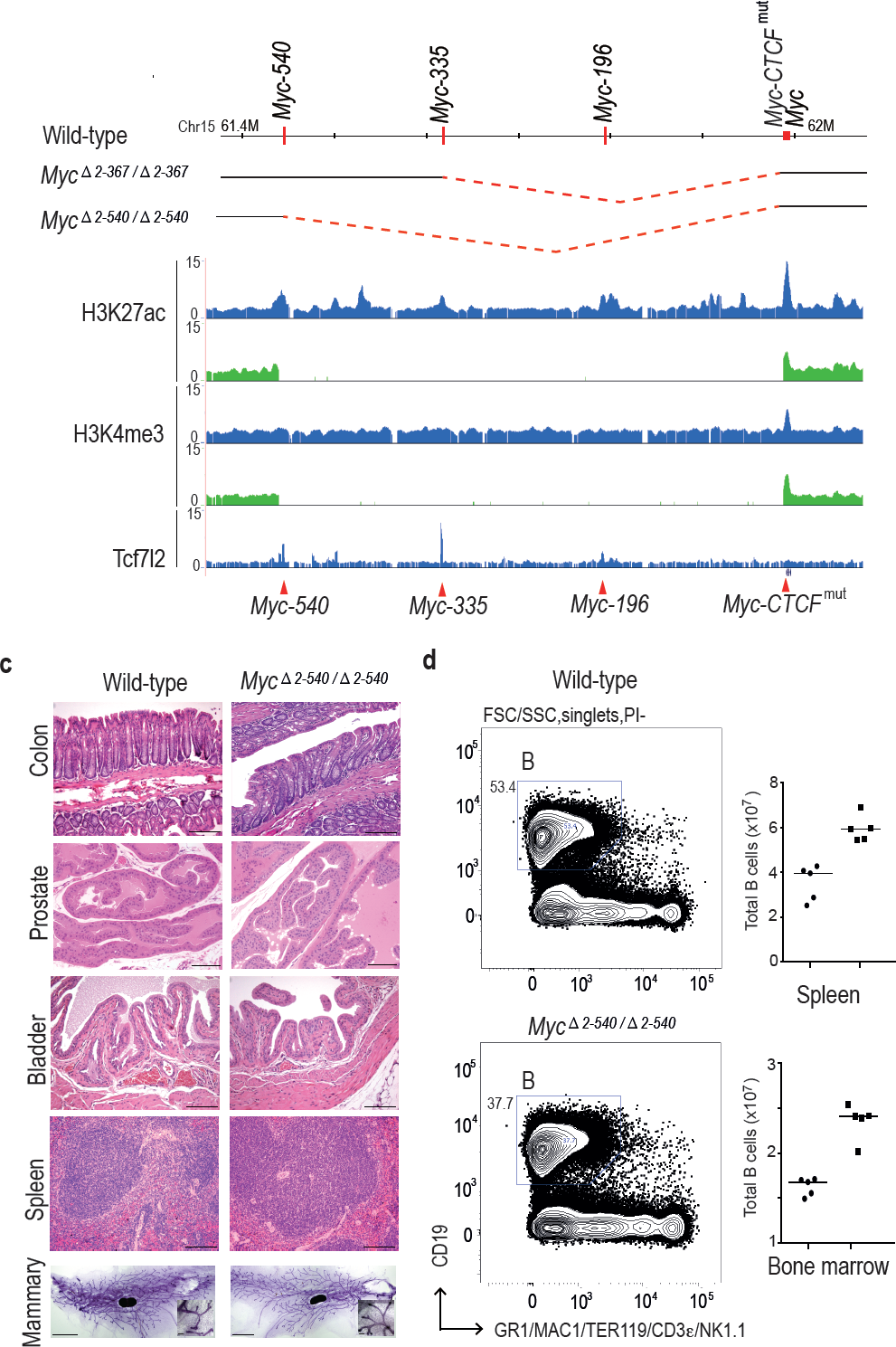
Deletion of the 8q24 super-enhancer region is well tolerated during normal development and homeostasis. (a) Schematic representation of the 365 kb and 538 kb deletions. (b) Myc^Δ^2-540/^Δ^2-540^^ deletion abolishes several active enhancer elements upstream of Myc as shown by ChIP-seq analysis of histone H3 lysine 27 acetylation (H3K27ac) and lysine 4 trimethylation (H3K4me3). Signal from Myc^Δ^2-540/^Δ^2-540^^ and wild-type mice are shown in green and blue, respectively. Red arrowheads and horizontal lines mark the different enhancer positions. (c) Haematoxylin/ Eosin stained sections of spleen, bladder, prostate, colon (Bar = 100 μm) and Carmine Alum stained whole mounts of mammary glands, Bars = 3 mm, 100 μm (inset) showing normal development and homeostasis of different organs in Myc^Δ^2-540/^Δ^2-540^^ mice. (d) Myc^Δ^2-540/^Δ^2-540^^ mice have a reduced number of B-cells compared to the wild-types. Left panel: FACS plots of a representative Myc^Δ^2-540/^Δ^2-540^^ and wild-type mouse spleen showing B-cell (B) population. Right panel: Scatter dot plot of total number of B cells in the spleen and bone marrow of wild-type (squares), n=5 and Myc^Δ^2-540/^Δ^2-540^^ (filled circles), n=5. Each point represents individual mouse. Line represents the median.

The larger deletion, Myc^Δ^2-540^^, could also be bred to homozygosity, and both males and females were viable. The viability of the mice is striking, given that the region deleted contains regions linked to chronic lymphocytic leukemia, and bladder, prostate, breast, and colon cancers. To characterize the mice further, we analyzed histology and Myc expression in the tissues where these tumors originate from. This analysis revealed normal morphology of mammary gland, spleen, bladder, prostate and colon in Myc^Δ^2-540/^Δ^2-540^^ mice (Fig. 2c).

### Loss of the super-enhancer region leads to tissue-specific changes in Myc expression

Although the Myc^Δ^2-540/^Δ^2-540^^ mice exhibited a normal phenotype, the expression of *Myc* was strongly decreased in colon, small intestine and prostate of these mice (Fig. 3a and not shown). Immunohistochemical analysis of Myc expression in intestine revealed strong decrease of nuclear staining, and loss of Myc expression from the transit amplifying cell compartment. However, expression of Myc was still detected at the base of the crypt in the region where the intestinal stem cells are known to reside (Fig. 3b). These results are consistent with the role of the deleted region in tumorigenesis of colon and prostate. To analyze the effect of decreased Myc expression on the proliferation in the transit amplifying compartment, we performed immunohistochemistry (IHC) for the proliferation marker Ki-67. Both the wild-type and Myc^Δ^2-540/^Δ^2-540^^ had similar proliferative activity in the intestinal crypts (Fig. 3b).

**Figure 3:**
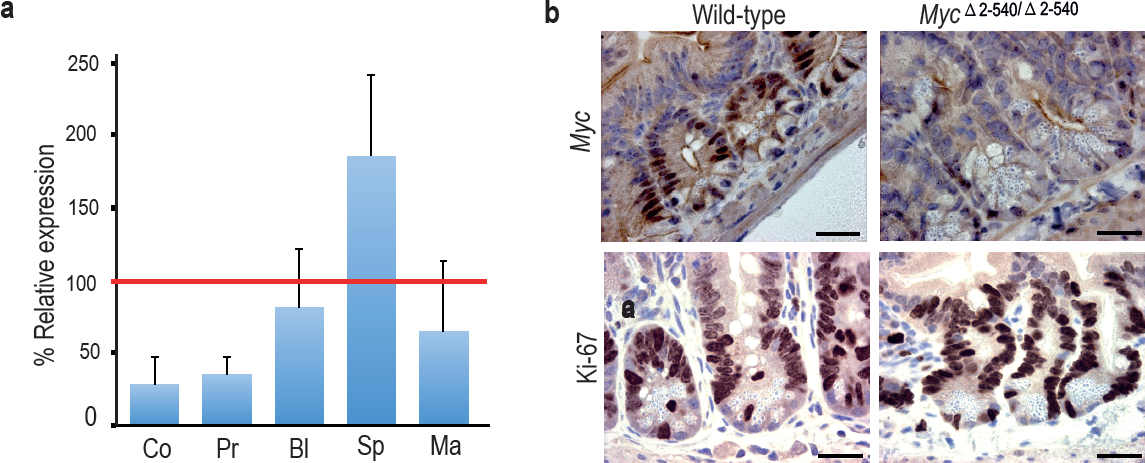
Tissue-specific effect of Myc^Δ^2-540/^Δ^2-540^^ deletion on Myc expression. (a) qPCR data showing the percentage of *Myc* expression in Myc^Δ^2-540/^Δ^2-540^^ relative to the wild-type in colon (Co) *n*=4, prostate (Pr) *n*=2, bladder (Bl) *n*=5, spleen (Sp) *n*=4 and mammary gland (Ma) *n*=3. Red line marks the expression level (100%) in wild-type mice. Error bars indicate one standard deviation. (b) Immunohistochemistry shows reduced expression of Myc protein in intestinal crypts of Myc^Δ^2-540/^Δ^2-540^^ mice without any significant effect on proliferation as indicated by Ki-67 immunostaining, Bar = 10 μm. Brown: IHC staining, Blue: Haematoxylin staining.

In contrast to colon and prostate, *Myc* expression was not markedly affected in the bladder, and was elevated in the spleen (Fig. 3a). To analyze the cellular composition of the spleen, we performed flow cytometric analysis of markers for hematopoietic stem cells and lymphoid lineage cells. Myc^Δ^2-540/^Δ^2-540^^ mice had a near normal hematopoietic compartment (Fig. 2d). The only observed difference was a small reduction of B cells in the Myc^Δ^2-540/^Δ^2-540^^ mice compared to the wild-type mice both in the spleen and the bone marrow. Although this is consistent with a role of this region in CLL development, this difference did not translate into a phenotypic consequence under normal laboratory conditions.

To compare the role of the 8q24 super-enhancer region in growth of cells *in vivo* and in cell culture, we isolated fibroblasts from the skin of adult Myc^Δ^2-540/^Δ^2-540^^ and wild-type mice. Based on presence of active histone marks, and undermethylation of focal elements (Fig. 4a and Supplementary Fig. S1), the super-enhancer region is active in fibroblasts from both humans and mice. However, the resident fibroblasts in the skin of Myc^Δ^2-540/^Δ^2-540^^ mice appeared normal as judged by Vimentin expression (Fig. 4b). Ki-67 staining (IHC) of skin sections showed comparable proliferation levels in wild-type and Myc^Δ^2-540/^Δ^2-540^^ mice (Fig. 4b). In contrast, most lines of fibroblasts (6 out of 7) isolated from Myc^Δ^2-540/^Δ^2-540^^ mice grew slower in culture compared to fibroblasts isolated from wild-type mice (Fig. 4c; *p*-value= 0.0256, Mann-Whitney one tailed test).

**Figure 4:**
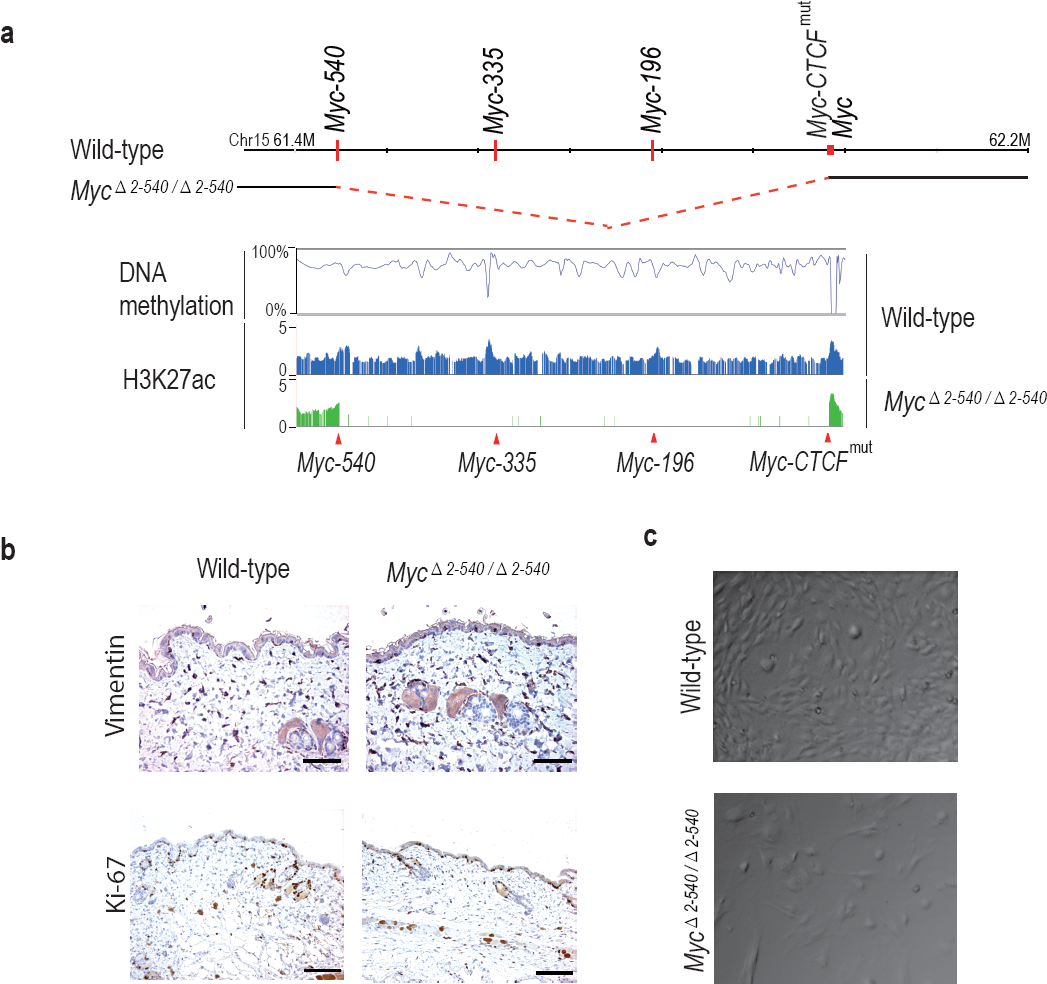
Myc^Δ^2-540/^Δ^2-540^^ deletion results in a proliferation defect of adult skin fibroblasts cultured *in vitro*. (a) The super-enhancer region deleted in the Myc^Δ^2-540/^Δ^2-540^^ has under methylated DNA as determined through bisulfite sequencing of the wild-type fibroblasts grown in culture. H3K27ac ChIP-seq shows the presence of active enhancer marks within this region in the wild-type fibroblasts whereas the Myc^Δ^2-540/^Δ^2-540^^ fibroblasts show a complete absence of the super-enhancer region. (b) Normal morphology and proliferation of resident fibroblasts in the mouse skin as determined by Vimentin IHC staining in both the wild-type and Myc^Δ^2-540/^Δ^2-540^^ mice, Bar = 50 μm. Brown: IHC staining, Blue: Haematoxylin staining (c) Representative phase contrast images of wild-type and Myc^Δ^2-540/^Δ^2-540^^ primary fibroblasts showing growth defect of Myc^Δ^2-540/^Δ^2-540^^ fibroblasts in culture.

### Deletion of the Myc super-enhancer region affects Myc target gene expression only in culture conditions

To understand the mechanism by which the deletion of the 8q24 super-enhancer region has a differential effect on growth during normal tissue homeostasis and growth under culture conditions, we subjected both the mouse tissues and cultured cells to RNA-seq analysis. Analysis of mouse tissues confirmed the changes in *Myc* expression observed by qPCR (Fig. 5a). Surprisingly, despite more than 80% decrease of *Myc* expression in the colon, very few genes were downregulated in the tissues, and none of the significantly altered genes were known Myc targets (Supplemental Table S1). These results suggest that expression of canonical Myc target genes is not sensitive to decreases in Myc protein level during normal tissue homeostasis. In contrast to the *in vivo* situation, where Myc is downregulated but key target genes are not affected, in cultured Myc^Δ^2-540/^Δ^2-540^^ fibroblasts that grew slowly in culture, the downregulation of *Myc* lead to a loss of expression of key target genes that drive cell growth and division. Upstream regulator analysis performed using Ingenuity Pathway Analysis revealed that the highest-ranked potential regulator for the identified gene set was Myc (Fig. 5b).

**Figure 5:**
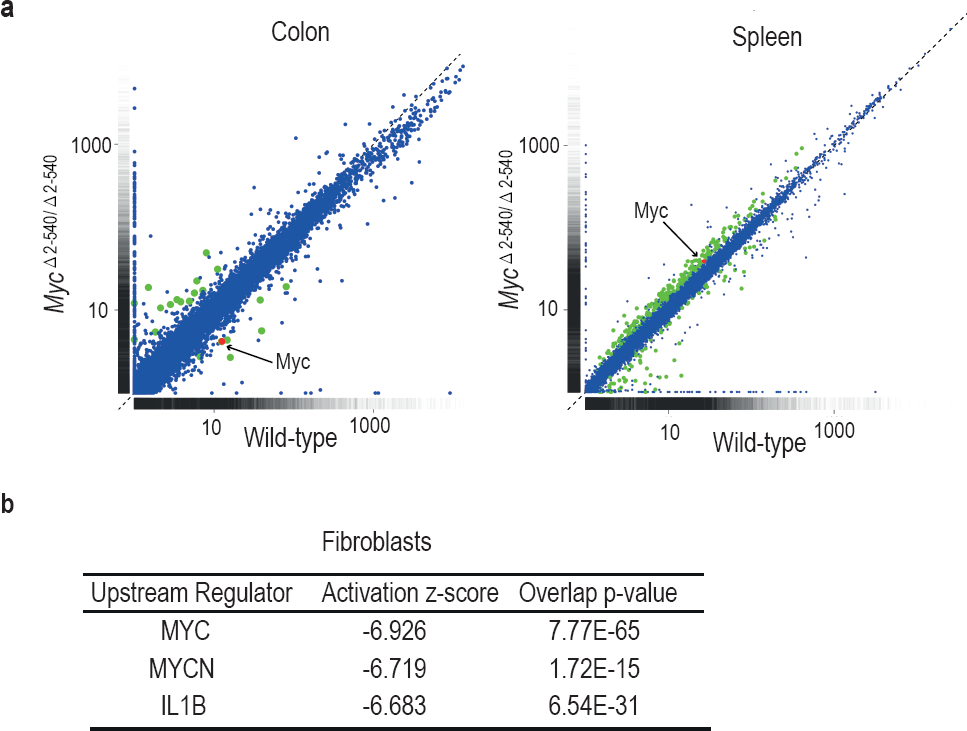
Differential effect of Myc^Δ^2-540/^Δ^2-540^^ deletion on Myc target gene expression. (a) Scatter plot comparing the average Fragments per kilobase of exons per million fragments mapped (FPKM) values of gene transcripts in colon and spleen of wild-type (*n*=4) and Myc^Δ^2-540/^Δ^2-540^^ (*n*=4) mice. Genes showing significant (q<0.05) differential expression are marked in red (Myc) or green (other genes). (b) Upstream regulator analysis of RNA-seq data shows that the highest ranked potential regulator affected in the slow growing Myc^Δ^2-540/^Δ^2-540^^ fibroblasts is Myc. The activation z-scores are to infer the activation states of predicted upstream regulators. The overlap *p*-values were calculated from all the regulator-targeted differential expression genes using Fisher’s Exact Test.

Measured by FPKM values, the cultured wild-type fibroblasts had higher *Myc* mRNA levels than normal tissues, whereas the cultured null fibroblasts had *Myc* levels that were comparable to or lower than those of normal wild-type tissues. The elevated *Myc* levels in cultured cells are caused by serum stimulation, as *Myc* mRNA levels are low in serum-starved fibroblasts and strongly induced by serum (Ref. (Dean et al. 1986) and our unpublished data). These results indicate that the 8q24 super-enhancer region is dispensable for normal tissue homeostasis under conditions where Myc activity is relatively low. However, the region is required for induction of Myc activity to levels that are high enough to drive the expression of Myc target genes above their basal levels during pathological growth.

### The Myc super-enhancer region is required for tumorigenesis in mice

We have shown earlier that deletion of a 1.8 kb MYC-335 enhancer sequence located at the 8q24 super-enhancer region is required for intestinal tumorigenesis in mice (Sur et al. 2012). As the super-enhancer region deleted in Myc^Δ^2-540/^Δ^2-540^^ mice carries risk also for leukemia, and prostate, breast, and bladder cancer, we tested the susceptibility of the Myc^Δ^2-540/^Δ^2-540^^ mice to carcinogen induced bladder and mammary tumorigenesis. The Myc^Δ^2-540/^Δ^2-540^^ mice were not resistant to N-Butyl-N(4-hydroxybutyl) nitrosamine (BBN) induced bladder tumors. Both wild-type (n=8) and Myc^Δ^2-540/^Δ^2-540^^ (n=8) mice developed urothelial changes ranging from hyperplasia to high grade invasive urothelial carcinoma after 5 months of BBN treatment. In contrast, comparison of median tumor-free survival times of wild-type and Myc^Δ^2-540/^Δ^2-540^^ mice exposed to mammary-tumor inducing dimethylbenz[a]anthracene/ medroxypregesterone (DMBA/MPA) regimen revealed that the Myc^Δ^2-540/^Δ^2-540^^ mice were partially resistant to mammary tumorigenesis (Fig. 6a). The median tumor-free survival time for the wild-type and Myc^Δ^2-540/^Δ^2-540^^ mice was 88 and >120 days, respectively. Together with our earlier findings, these results indicate that loss of the 8q24 super-enhancer region makes mice resistant to both genetically and chemically induced tumors.

**Figure 6:**
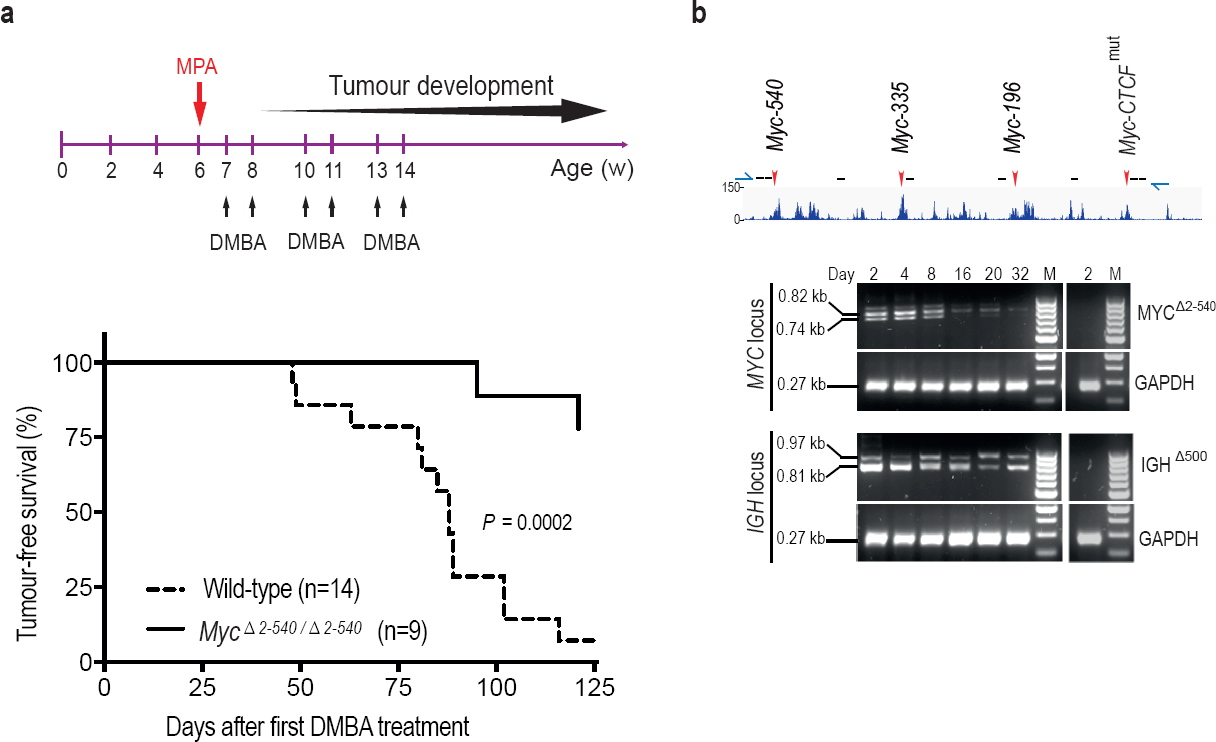
*Myc* -2 to -540 kb genomic region is required for the growth of cancers *in vivo* and cancer cells *in vitro*. (a) Tumor-free survival plots showing resistance of Myc^Δ^2-540/^Δ^2-540^^ mice to development of DMBA/MPA induced mammary tumors. p-value =0.0002 (Mantel-Cox Log-rank test). (b) Crispr-Cas9 mediated deletion of region corresponding to *Myc*^Δ^2-540/^Δ^2-540 region^^ in human GP5d colon cancer cells, results in a loss of the edited cells over time. Top panel shows the active enhancer elements in GP5d cells within this region as determined by ChIP-seq analysis of histone H3 lysine 27 acetylation (H3K27ac). The sites of sgRNAs (black lines) and genotyping primers (green arrows) used are indicated (not to scale). Red arrows mark the enhancer regions used in this study. Bottom panel shows the PCR-genotyping of the MYC locus and the control IGH locus showing the specific loss of the edited MYC locus over time. GAPDH was used as internal control. The right panel in each set shows absence of any deletion in the non-transfected cells (day 2). 100 bp ladder DNA molecular weight marker is shown (M)

We further tested the requirement of this region for the proliferation of cancer cell lines in cultures. We found that the corresponding region (hg19: chr8:128226490-128746456) was also required for GP5d colon cancer cell growth, as indicated by progressive loss of cells bearing a CRISPR/Cas9 induced deletion of the region during co-culture with unedited cells in the population (Fig. 6b).

## Discussion

The region around the *MYC* gene carries inherited risk towards multiple major forms of cancer. On aggregate, this region contributes more to inherited cancer than any other locus in the human genome. The risk alleles for different cancer types are located in multiple distinct linkage disequilibrium blocks, indicating that different variants contribute to different cancer types. Several of these regions containing risk variants have been implicated in regulation of MYC expression (Hallikas et al. 2006; Sur et al. 2012; Herranz et al. 2014; Uslu et al. 2014), suggesting that a large number of enhancers within this region can drive tumorigenesis. Some of the identified elements have also been shown to have roles in normal development (Herranz et al. 2014; Uslu et al. 2014).

To study the role of the 8q24 region more systematically, we have in this work deleted several individual elements, and also analyzed the effect of larger deletions on normal development and carcinogenesis in mice. Our analysis of mice lacking a 538 kb region upstream of the *Myc* gene suggests that enhancer elements within this region cooperatively enhance Myc expression. Deletion of individual enhancers in this region has only a weak (Sur et al. 2012) or no effect on Myc expression in the mouse intestine in contrast to the deletion of the entire super-enhancer region, which leads to severe decrease in Myc expression in multiple tissues.

Myc deficient mouse embryos die due to placental defect at E9.5. When Myc is deleted only in the epiblast, the embryos grow normally and survive until E11.5, when they die due to defects in hematopoiesis (Dubois et al. 2008). None of these defects are observed in mice homozygous for the super-enhancer deletion showing that the 8q24 super-enhancer region is dispensable for Myc function both in the placenta and during early hematopoiesis.

Despite decreased levels of Myc in multiple adult tissues, the mice lacking the super-enhancer region display normal tissue morphology in all the tissues we investigated, and do not have marked defects in cell proliferation. The mice are, however, resistant to DMBA-induced mammary tumors, indicating that this region is important for tumorigenesis also in mice. These results indicate that despite the central role of this region in tumorigenesis (Sur et al. 2012; Loven et al. 2013), it is dispensable for normal tissue development and homeostasis under laboratory conditions. Whereas this result may appear very surprising, it is consistent with the original identification of this region using genome-wide association studies (GWAS). GWAS has a high power to identify common variants, and most variants that are common have only a limited effect on physiological functions. This is because a variant that has strong positive or negative effect is rapidly fixed or lost, respectively. Thus, GWAS are specifically biased to find variants that have a relatively large effect on disease, but a small effect on fitness.

Most genes in mammals do not have haploinsufficient phenotypes. Buffering could be due to mechanisms that maintain constant expression level irrespective of gene dose. However, a simpler buffering mechanism involves either expressing a gene at a very low level where it has no effect, or at a high level where it can contribute its functions even if its expression level is decreased due to transcriptional noise or loss-of-function of one allele. A similar two state mechanism where physiological TF activity levels in the relevant cell types are either too low to drive any target genes (off state), or high enough to activate all important targets (on state) could also mechanistically explain why most heterozygous null mutations of TF genes have no apparent phenotype. Our analysis of the role of Myc in normal colon is consistent with such a simple buffering model (Fig. 7). Under normal physiological conditions, the system is in the off state, and a basal level of expression of the Myc target genes is maintained by a Myc-independent mechanism. The genes are thus only sensitive to an increase in Myc levels. Consistently, an 80% decrease of *Myc* mRNA expression does not lead to a proliferation defect, or major changes in expression of known Myc target genes. In contrast, in tumors the system is locked to an on state, where Myc targets are driven to a maximal level by Myc, and the targets are now only sensitive to a decrease in Myc activity (Fig. 7).

**Figure 7:**
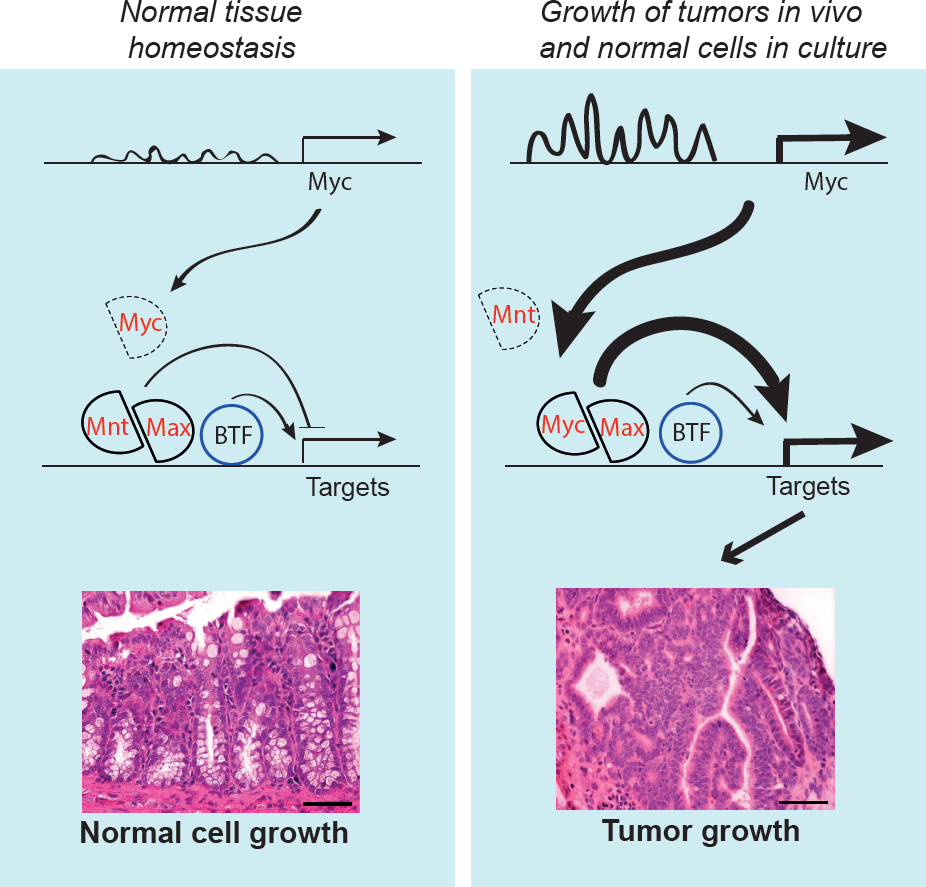
Model showing the activity of the *Myc* super-enhancer during normal homeostasis (left) and cancer (right). During normal tissue homeostasis (left), *Myc* enhancers are not strongly active, and Myc activity is relatively low. The Myc expression level is insufficient to recruit enough Max proteins to Myc/Max heterodimers to drive strong induction of the Myc target genes, which instead remain under the control of basal transcription factors (BTF). In cancer cells or cells grown in culture (right), upstream regulators such as Tcf7l2 and β-catenin activate the *Myc* super-enhancers, driving high levels of Myc expression. This leads to the formation of Myc/Max heterodimers that strongly activate transcription of Myc target genes, which drive cancer cell growth. The Myc super-enhancer region, and induction of the Myc target genes are not required for normal homeostasis, indicating that they are promising targets for antineoplastic therapies.

The requirement of MYC in tumor cells appears absolute. In transgenic animal models, overexpression of Myc leads to deregulated proliferation and tumor development in multiple tissues(Felsher and Bishop 1999; Pelengaris et al. 1999; D’Cruz et al. 2001; Jain et al. 2002; Shachaf et al. 2004). Furthermore, inhibition of MYC almost invariably causes growth arrest of cancer cells both in culture and *in vivo* (Soucek et al. 2002; Soucek et al. 2004; Hart et al. 2014). Despite the importance of Myc for cancer growth, it appears that the role of Myc in controlling growth during normal development is limited. In the adult tissues, Myc is expressed in rapidly proliferating compartments of the body like the intestinal crypts and skin. Deletion of *Myc* in these compartments does not result in prominent proliferation defects (Wilson et al. 2004; Baena et al. 2005; Bettess et al. 2005; Muncan et al. 2006). Although there is still controversy regarding Myc requirement for the intestinal homeostasis, in the skin Myc is dispensable under normal adult proliferation and homeostasis *in vivo* (Oskarsson et al. 2006). It is however required for Ras mediated tumorigenesis and growth of fibroblasts and keratinocytes *in vitro* (Mateyak et al. 1997; Oskarsson et al. 2006). Taken together, these results suggest that MYC is required for pathological proliferation, but is less important and in many cases dispensable for normal homeostasis of tissues. Our results are consistent with these observations. However, prior to our study, the mechanism behind this differential activity was unclear. In addition, it has not been clear whether Myc expression is regulated via similar mechanisms in cultured normal cells and cancer cells. Our results have uncovered striking mechanistic similarities between growth of normal cells in culture, and growth of cancer cells *in vivo*. They also suggest that many potential cancer drugs may have been inadvertently discarded due to their negative effects on growth of normal cells in culture.

Our results show that the MYC super-enhancer that carries multi-cancer susceptibility in humans contributes to the formation of multiple tumor types also in mice. Despite its role in tumor formation, it is dispensable for normal development and homeostasis. Loss of the super-enhancer leads to low Myc expression, but the lowered expression does not translate to changes in expression of Myc target genes during normal development. Thus, the MYC/MAX/MNT system (Grandori et al. 2000) that drives cell growth and proliferation is robustly set to an off state during normal homeostasis, whereas in cancer, the system is locked to a pathological on state. This also explains how physiological growth control can be robust to small perturbations and transcriptional noise. Taken together, our results reveal an important difference between the transcriptional states of normal and cancer cells, and suggest that therapeutic interventions that decrease the activity of the Myc super-enhancer region would be well tolerated.

## Materials and methods

### Mouse strains

We generated cKO Myc-196 and cKO Myc-540 strains with loxP sites flanking the regions chr15:61445326-61447611 and chr15:61789274-61791107, respectively (Taconic). These mice were crossed to EIIa-cre mouse strain (Jackson Laboratory) to generate mice with enhancer deletions. Myc-CTCF^mut^ mouse strain was generated by mutating the CTCF-binding site at chr15:61983375-61983647 TGGCCAGTAGAGGGCAC to TGGAACGTCTTGAATGC. In order to generate large deletions at the Myc locus (Myc^Δ^2-367^^ and Myc^Δ^2-540^^) Myc-367 and Myc-540 were crossed to Myc-CTCF^mut^ that were also heterozygous for the Rosa26-Cre (Taconic). The Myc-540, Myc-196 and Myc-CTCF^mut^ carry one lox P site at the respective loci (chr15:61445326, chr15:61618287 and chr15:61983375). The loxP site on chr15:61983375 is located immediately 5’ of the mutant CTCF binding site. We obtained compound heterozygotes carrying the chr15:61445326 or the chr15:61618287 loxP site together with the loxP site on chr15:61983375 and the Rosa26 Cre. The compound heterozygotes were screened by PCR for the interallelic recombination and the resultant deletion and duplication of the intervening sequence. Mice mosaic for the deletion and duplication were backcrossed to the C57Bl/6 mice in order to segregate the chromosomes carrying the deletion. The F1 heterozygotes were intercrossed to generate mice with homozygous large deletions. Myc-335 strain has been previously described (Sur et al. 2012). All mice used in the study were on a C57Bl/6 genetic background. Mouse experiments were conducted in accordance with the local ethical guidelines. The sequences of the different primer pairs used for genotypings are given in Supplemental Table S2.

### Mammary gland whole mount analysis

Inguinal mammary glands were removed from 8 week old virgin females and spread on glass slides. These were fixed for 4 hours in Carnoy’s fixative and subsequently stained O/N with Carmine Alum. The whole mounts were rinsed and dehydrated through increasing series of ethanol and cleared in xylene before mounting with the pertex mounting medium.

### Quantitative PCR analysis

qPCR was performed as described previously (Sur et al. 2012). Essentially, total RNA was isolated from whole tissue by homogenizing in RNA Bee reagent (ambios AMS Biotechnology) followed by RNA isolation using Qiagen’s RNA MinElute kit according to manufacturers’ protocols. 0.5-1 µg of total RNA was reverse transcribed using high capacity reverse transcription kit in a 20 µl reaction (Applied Biosystems). Quantitative PCR in triplicates was performed using the SYBR select master mix (Applied Biosystems) on the LightCycler 480 instrument (Roche). For normalization, mouse β -actin transcripts were used as internal controls. Following primer pairs were used for quantitative PCR analysis.

Myc-Fw: 5’-GGGGCTTTGCCTCCGAGCCT-3’, Myc-Rev: 5’-TGAGGGGCATC GTCGTGGCT-3’, β-actin-Fw: 5’CTGTCGAGTCGCGTCCACCCG-3’, β-actin-Rev: 5’-CATGCCGGAGCCGTTGTCGAC-3’.

### RNA-sequencing

NEBNext Ultra Directional RNA library Prep kit (NEB) was used for preparing the samples for RNA-seq together with the NEBNext Poly(A) mRNA magnetic isolation module (NEB) according to manufacturers protocol. In the case of tissues 1-2 μg and for cultured fibroblasts 200 ng of total RNA was used as starting material. For library preparation, adapters and index primers from NEBNext Multiplex Oligos for Illumina kit were used. The RNA-seq library was sequenced on a HiSeq2000 (Illumina). Sequencing reads were mapped to the mouse reference genome (NCBI37/mm9) using Tophat2 (version 2.0.13)(Kim et al. 2013). Cuffdiff (version 2.2.1) was used for differential gene expression analysis and for graphical representation, CummeRbund package (version 2.8.2)(Trapnell et al. 2012) was used. The upstream regulator analysis was performed on all the significant differentially expressed genes (Cuffdiff q-value < 0.05) using QIAGEN’s Ingenuity Pathway Analysis (IPA, QIAGEN Redwood city, www.qiagen.com/ingenuity; version 24718999, updated 2015-09-14).

### ChIP-seq

ChIP-seq was performed as described in (Sur et al. 2012; Yan et al. 2013) with the following modifications: Adult 8-10 week old mice were euthanized and colon was removed, rinsed with cold PBS and cut into fine pieces. Tissue was crosslinked with 1.5% formaldehyde and cultured cells were crosslinked with 1% formaldehyde for 10 minutes at room temperature and quenched with 0.33M Glycine. Sequences were mapped to the mouse reference genome (NCBI37/mm9) using Burrows-Wheeler Alignment tool (bwa) (version 0.6.2)(Li and Durbin 2009) with default parameters. All antibodies used in ChIP-seq experiments were ChIP-grade. In each experiment a non-specific IgG was used as control. Following antibodies were used for ChIP-seq experiments: rabbit anti-H3 lysine 27 acetylation (H3K27ac) (abcam, ab4729), mouse anti-H3 lysine 4 trimethylation (H3K4me3) (abcam, ab1012), rabbit anti-Rad21 (Santa Cruz, sc-98784), goat anti-CTCF (Santa Cruz, sc-15914X), rabbit anti-SMC1A (Bethyl Laboratories, A300-055A), rabbit IgG (Santa Cruz, sc-2027), mouse IgG (Santa Cruz, sc-2025), goat IgG (Santa Cruz, sc-2028). ChIPseq data for Tcf7l2 was used from ENA accession number PRJEB3354 (Sur et al. 2012) and for GP5d cells from ENA accession number PRJEB1429 (Yan et al. 2013). For visualization, ChIP-seq read depth data were average smoothed across windows of 10 pixels (H3K27ac and H3K4me3) or 5 pixels (Tcf7l2) in UCSC Genome Browser or alternatively visualized in Integrative Genomics Viewer (IGV, version 2.3)

### Bisulfite sequencing

Genomic DNA was isolated using Qiagen’s Blood & Tissue Genomic DNA extraction kit. Around 1 µg of wild-type and 250 ng of Myc^Δ^2-540/^Δ^2-540^^ null fibroblast genomic DNA was sonicated to 300bp fragments using Covaris S220 sonicator. Subsequent to end polishing and A base addition, cytosine methylated paired end adapters (Integrated DNA technologies) were ligated to the DNA fragments. The adapter sequences are as follows

5’P-GATCGGAAGAGCGGTTCAGCAGGAATGCCGAG

5’ACACTCTTTCCCTACACGACGCTCTTCCGATCT

After adapter ligation 300-600 bp fragments were size-selected on a 2% agarose gel. Bisulfite-conversion was carried out using ZYMO EZ DNA Methylation-Gold kit (cat. no. D5005). PCR amplification with 12 and 18 cycles was carried out to prepare libraries from the wild-type and Myc^Δ^2-540/^Δ^2-540^^ null mouse fibroblasts, respectively. The primer pair used for PCR amplification was as follows

PE PCR Primer P1

5’AATGATACGGCGACCACCGAGATCTACACTCTTTCCCTACACGACGCTCT TCCGATCT

PE PCR Primer P2:

5’CAAGCAGAAGACGGCATACGAGATCGGTCTCGGCATTCCTGCTGAACCG CTCTTCCGATCT

The final library was size-selected for 250-300 bp fragments on a 2% agarose gel and 150 bp sequenced from both ends on two lanes of a HiSeq 4000 (Illumina). Raw sequencing reads were quality and adapter trimmed with cutadapt version 1.8.1 in Trim Galore version 0.4.0. Trimming of low-quality ends was done using Phred score cutoff 30. In addition, all reads were trimmed by 2 bp from their 3’ end. Adapter trimming was performed using the first 13 bp of the standard Illumina paired-end adapters with stringency overlap 2 and error rate 0.1. Read alignment was performed against mouse genome mm9 with Bismark (version v0.14.3) (Krueger and Andrews 2011) and Bowtie 2 (version 2.2.4)(Langmead and Salzberg 2012). Duplicates were removed using the Bismark deduplicate function. Extraction of methylation calls was done with Bismark methylation extractor discarding first 10 bp of both reads and reading methylation calls of overlapping parts of the paired reads from the first read (-no_overlap parameter). Genomic sites with the coverage of at least 10 reads were considered and methylation ratios smoothed with loess method across 49 bp windows. All sequencing data will be uploaded to European Nucleotide Archive (ENA, EMBL-EBI) under accession number PRJEB11397.

### Immunohistochemistry and flow cytometry

Five micron paraffin embedded tissue sections were processed for immunohistochemistry as previously described (Sur et al. 2012). Rabbit polyclonal anti-Myc (Santa Cruz) (1:500), Rabbit monoclonal anti Ki-67 (abcam) (1:200), Goat polyclonal anti-Vimentin (Santa Cruz) (1:500), biotinylated goat anti-Rabbit IgG and biotinylated rabbit anti-Goat IgG (Vector Laboratories) (1:350) antibodies were used. For flow cytometry, single cell suspensions of spleen cells were stained with Fc-block (CD16/CD32 clone 93) and subsequently with CD19 (1D3), TER119 (TER119), CD3ε (145-2C11), NK1.1(PK136), GR1/LY6G (1A8) (BD). Dead cells were visualized using Propidium iodide. Samples were analyzed using a BD LSRFortessa instrument.

### Isolation and culture of mouse primary fibroblasts

Fibroblasts were isolated from adult mice by dissecting the skin to ~ 1 mm^3^ pieces, and allowing the pieces to adhere to cell culture plates, followed by addition of DMEM medium supplemented with 10% FCS and antibiotics. The fibroblasts were allowed to migrate out from the explants, after which the cells were collected by trypsinization and passaged in the same media for 1-3 passages. For growth assays, 2x10^3^ cells were plated per well in 96 well plates. Cells were trypsinized and counted using hemocytometer at respective time points.

### Tumor induction

Mammary tumors: Six week-old female mice were implanted s.c. with medroxypregesterone acetate (MPA) pellets (50 mg with a 90 days release period from Innovative Research of America). Subsequently 100 μl of 10 mg/ml dimethylbenz[a]anthracene (DMBA)/oil solution (Sigma) was administered via gavage at 7,8, 10,11, 13 and 14 weeks of age. Mice were checked twice a week for development of palpable tumors. Detection of palpable tumors was taken as the end point for tumor-free survival analysis.

Bladder tumors: Ten week-old male mice were administered 0.1% N-Butyl-N-(4-hydroxybutyl) nitrosamine (BBN) (Sigma) in drinking water for 5 months. At the end of the treatment the mice were sacrificed and the bladders scored for tumor development.

### CRISPR-Cas9 mediated deletion of Super-enhancer in GP5d cell line

CRISPR-Cas9 mediated deletion of *MYC* super-enhancer region on chromosome 8q24 (GRCh37/hg19 chr8: 128226403-128746490) and Immunoglobulin Heavy (IGH) gene locus on chromosome 14q32.33 (GRCh37/hg19 chr14: 106527004-107035452) were carried out in GP5d (Sigma, 95090715) colon cancer cell line stably expressing Cas9 protein. A lentiviral plasmid containing Cas9 fused via a self-cleaving 2A peptide to a blasticidin resistance gene, was packaged into lentiviral particles using the packaging plasmids psPAX2 (a gift from Didier Trono, Addgene plasmid # 12260) and pCMV-VSV-G (a gift from Robert Weinberg (Addgene plasmid # 8454). The virus was used to transduce GP5d colon cancer cells. 48 hours after transduction, GP5d cells expressing Cas9 (GP5d-Cas9) were selected in 5µg/ml Blasticidin (Thermo Fisher Scientific Inc., Cat. no. A1113903). The single guide RNA (sgRNA’s) were designed (http://www.broadinstitute.org/rnai/public/analysis-tools/sgrna-design) to span the entire *MYC* super-enhancer region and *IGH* locus (Fig. 6), respectively (Eurofins MWG Operon). sgRNAs were cloned into an sgRNA Cloning Vector (Addgene Plasmid #41824) using Gibson assembly master mix (NEBuilder HiFi DNA assembly Master Mix, Cat no. E2621S). GP5d-Cas9 (2 x 10^6^) cells were transfected (using FuGENE HD Transfection Reagent, Cat.no E2312) with 10 µg of eight pooled equimolar sgRNA constructs. Post transfection half of the cultured cells were collected for PCR genotyping, while the other half was re-plated for culturing. Cells were collected at day 2, 4 and subsequently every 4^th^ day till day 32. DNA from cells was extracted (using DNeasy Blood & Tissue Kit, Qiagen Cat. no. 69506) and genotyped with 300 ng of DNA at following conditions - Initial denaturation of 95°C for 5 mins; denaturation of 98°C for 15 sec, annealing at 60°C for 30 sec, extension at 72°C for 30 sec (30 cycles for *MYC* super-enhancer and 35 cycles for *IGH* gene locus deletion genotyping); final extension at 72°C, 5 min. The sequences of the different guide RNAs and primer pairs used for PCR genotyping of the deletions are given in supplemental table S2. GP5d cells were cultured in DMEM supplemented with 10% FBS and antibiotics. The cell line was mycoplasma free.

## Acknowledgements

We thank Drs. Minna Taipale and Bernhard Schmierer for critical review of the manuscript, and Maria Hoh, Lijuan Hu and Agneta Andersson for technical assistance. We also thank Prof. Björn Rozell and the Morphological phenotype analysis core facility at KI for help with the sectioning and analysis of tissues. This work was supported by the Knut and Alice Wallenberg Foundation, Center for Innovative Medicine at Karolinska Institutet, and the EU FP7 collaborative project SYSCOL.

